# Multi-landmark alignment of genomic signals reveals conserved expression patterns across transcription start sites

**DOI:** 10.1101/2021.09.02.458606

**Authors:** Jose M. G. Vilar, Leonor Saiz

## Abstract

The prevalent one-dimensional alignment of genomic signals to a reference landmark is a cornerstone of current methods to study transcription and its DNA-dependent processes but it is prone to mask potential relations among multiple DNA elements. We developed a systematic approach to align genomic signals to multiple locations simultaneously by expanding the dimensionality of the genomic-coordinate space. We analyzed transcription in human and uncovered a complex dependence on the relative position of neighboring transcription start sites (TSSs) that is consistently conserved among cell types. The dependence ranges from enhancement to suppression of transcription depending on the relative distances to the TSSs, their intragenic position, and the transcriptional activity of the gene. Our results reveal a conserved hierarchy of alternative TSS usage within a previously unrecognized level of genomic organization and provide a general methodology to analyze complex functional relationships among multiple types of DNA elements.

## Background

Genomic signals encapsulate highly detailed quantitative information up to the nucleotide level^1^ on key aspects of DNA transcription, the subsequent RNA processing, and multiple DNA-dependent processes, including DNA methylation^2^, transcription factor binding^3^, and CRISPR-Cas9 efficiency^4^. At the core of interpreting this information, there are specific genomic locations, or genomic landmarks, such as TSSs, transcription factor binding sites, RNA splice junctions, or the midpoint of a DNA extended region^5^. These landmarks provide anchoring points to summarize general trends and characterize different types of DNA regions.

The prototypical approaches to analyze these data start with the alignment of the signals to a landmark along a one-dimensional coordinate for subsequent processing. In mathematical terms, the alignment of a genomic signal *g*(*z*) along the coordinate *z* to a landmark with position denoted by *z*_*U*_ leads to a relative coordinate *x* = *z* − *z*_*U*_ and an aligned signal *g*(*x* + *z*_*U*_). The most widely used type of processing is the aggregation of alignments for multiple positions *z*_*U*_ of the landmark, which leads to an average signal 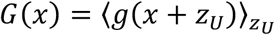. This approach has provided general information as diverse as the sharp dependence of CRISPRi/a activity on both the proximity of a TSS and nucleosome occupancy^6,7^; how the directionality of promoters reflects on the asymmetry of DNA accessibility and histone methylation signals around TSSs^8^; and the enrichment or depletion of single nucleotide variation occurrence around multiple landmarks in the genomes of human populations^9^. To capture the inherent heterogeneity, aligned signals are often structured into heatmaps^10^, which can be sorted and clustered according to specific parameters^11^ and can be incorporated into automated machine-learning pipelines^12^. This type of one-dimensional alignments is also the usual approach to link genomic signals with the results of methodologies, such as chromosome conformation capture techniques^13,14^, that map the three-dimensional DNA looping^15,16^ interactions between distal DNA elements.

The alignment with respect to a single position, however, is frequently ambiguous because regulatory regions often involve multiple relevant landmarks^17,18^. The presence of a landmark, such as a TSS, can often affect the functioning of another one and, in general, multiple landmarks can affect each other’s function. To analyze functional relationships among multiple types of DNA elements, we develop a method to consider multiple landmarks at the same level (Figure 1). The main idea is to align the signal to multiple locations through the expansion of the dimensionality of the genomic-coordinate space by considering relative coordinates from the different landmarks.

**Figure 1:**
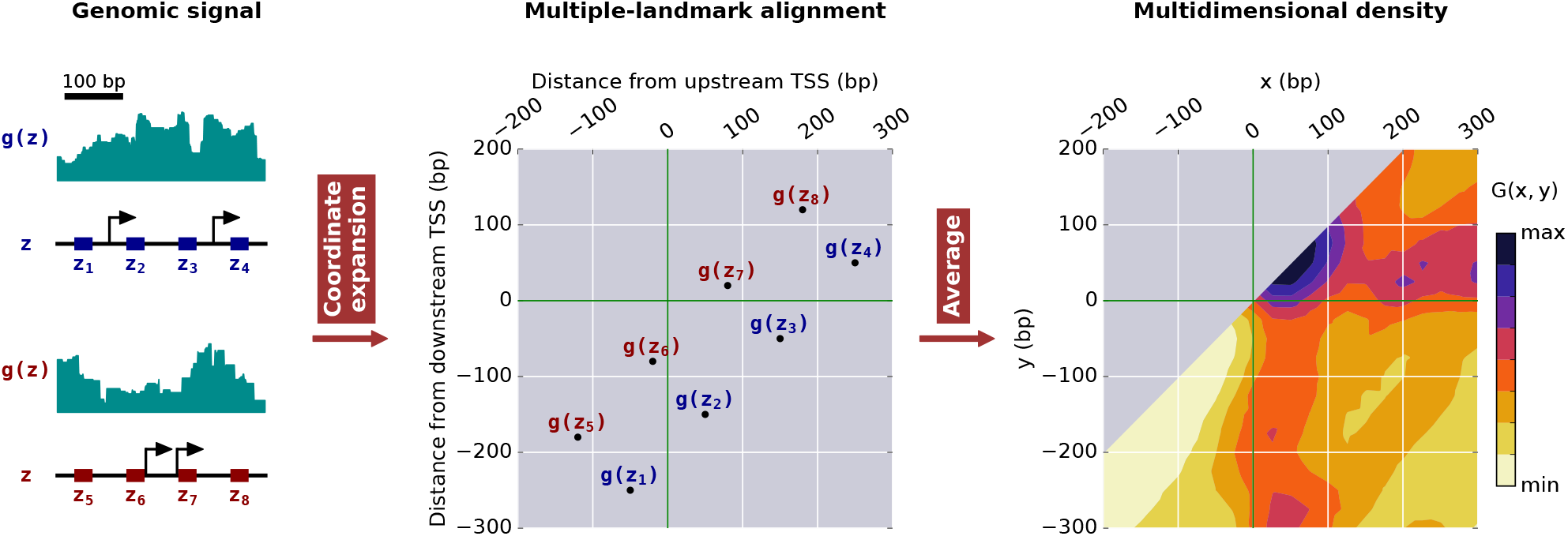
Constructing multidimensional representations of genomic signals. Starting with a genomic signal *g*(*z*) along the genomic coordinate *z*, we perform a coordinate expansion using multiple landmarks, such as TSSs (depicted by black arrows), to obtain a multiple-landmark alignment of the signal. For pairs of landmarks, genomic locations in the neighborhood of two landmarks, such as those in the intervals *z*_1_ − *z*_4_ and *z*_5_ − *z*_8_, are mapped into a two-dimensional representation with respect to the distances from each of the landmarks. Taking the average of *g*(*z*) in the expanded space for all the relevant pairs of landmarks provides a multidimensional signal density, depicted by *G*(*x*, *y*) in two dimensions.

## Results

### Simultaneous alignment to multiple positions

To consider genomic signals in two dimensions, we expand the genomic coordinate *z* with respect to the positions of the upstream, *z*_*U*_, and downstream, *z*_*D*_, landmarks into *x* = *z* − *z*_*U*_ and *y* = *z* − *z*_*D*_. For each pair of landmarks, the fact that *x* and *y* correspond to the same genomic coordinate leads to a line in the *x*, *y*-plane defined by *y* + *z*_*D*_ = *x* + *z*_*U*_. The signal along this line in the two-dimensional space can be described mathematically by 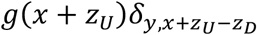, where *δ_i,j_* represents the Kronecker delta function, which is one if *i* = *j* and zero otherwise. This description allows the efficient computation of the two-dimensional average signal density, 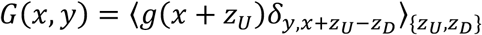, over pairs of landmarks {*z*_*U*_, *z*_*D*_} and a sliding window around (*x*, *y*). The use of the Kronecker delta function is also useful because it allows the straightforward extension of the methodology to multiple dimensions. For instance, in the case of three locations, the aligned three-dimensional signal is given by 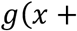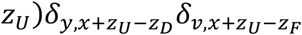, where *ν* = *z* − *z*_*F*_ is the relative position associated with the landmark with position *z*_*F*_.

### Complex dependence of transcription on multiple TSSs

The resulting multidimensional signal density provides a precise general description to analyze any function of a genomic coordinate in terms of the distances from multiple genomic landmarks. We use this approach to study the dependence of transcription, as reported by RNA sequencing (RNA-seq), on pairs of consecutive TSSs. Specifically, we focus on how transcription in human at a given genomic location depends on the relative positions of two TSSs, including how the presence of a TSS correlates with transcription at another TSS. The transcription of mammalian genomes^19,20^, with an average of four TSSs per gene^21^, is particularly relevant because the arrangement of TSSs according to different positional patterns, such as those in focused or dispersed promoters, is associated with different types of transcriptional programs^22^. TSSs locations were obtained from the comprehensive gene annotation on the reference chromosomes of Gencode V19. By considering the comprehensive set of annotated TSSs rather than only the ones expressed in each particular cell type, we could also investigate the factors that correlate with alternative TSSs expression.

As a representative case, we consider explicitly K562 human myeloid leukemia cells for the first and second TSSs (Figures 2A and S1A) and second and third TSSs (Figure 2B and S1B) of each protein-coding gene. Here, *TSSs are ordered according to their genomic position, starting the enumeration from the most upstream TSS*. The two-dimensional RNA-seq signal density *G*(*x*, *y*) reveals a strong dependence on the relative position of pairs of TSSs. There are dominant trends, such as a high transcriptional signal density downstream of the TSSs and the suppression of the signal upstream of a TSS.

**Figure 2:**
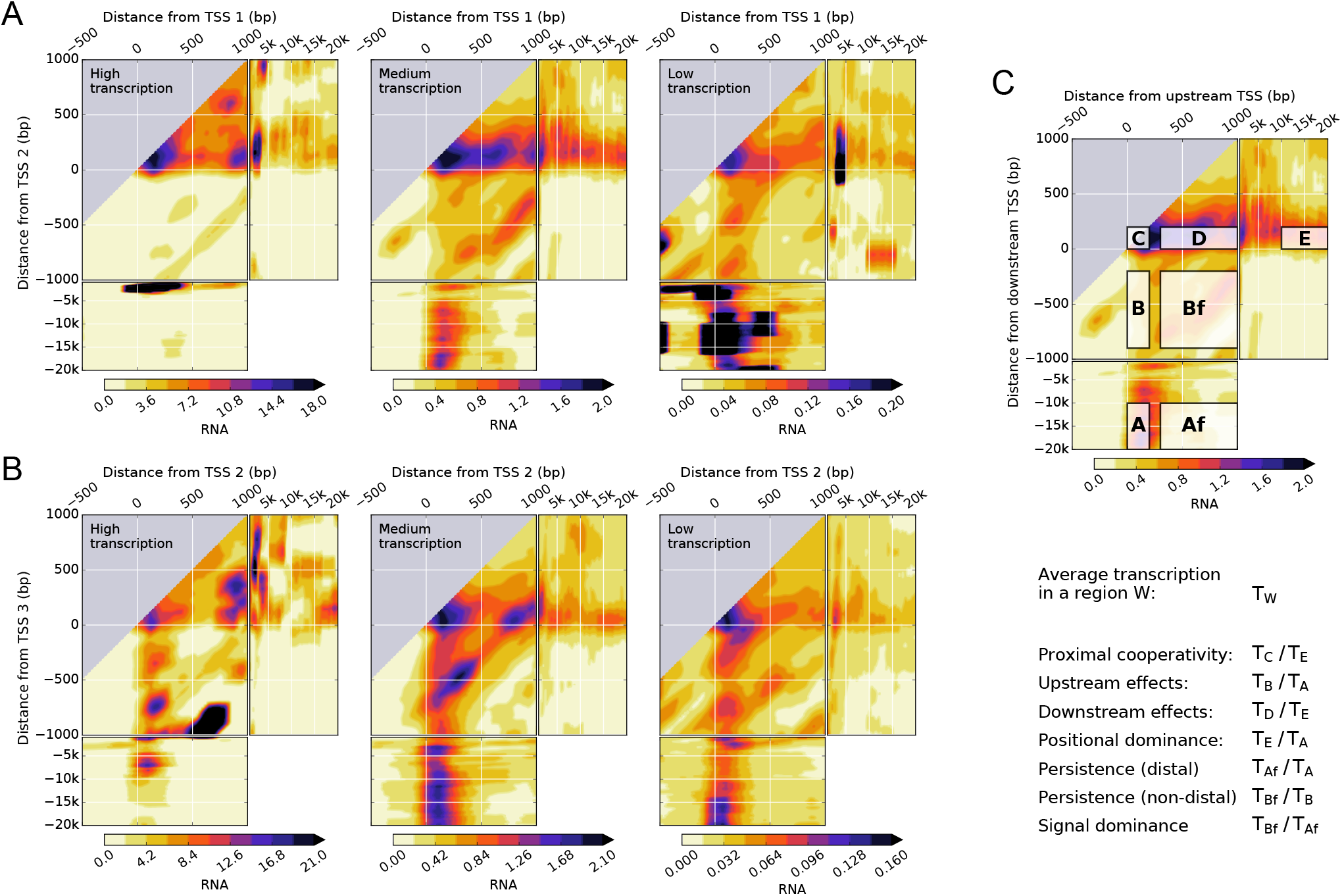
Transcription in K562 leukemia cell lines shows a complex dependence on the distance from pairs of TSSs, their intragenic position, and the transcriptional activity of the gene. (**A**, **B**), two-dimensional density of normalized RNA-seq signal for pairs of the first (TSS 1) and second (TSS 2) TSSs (**A**) and the second (TSS 2) and third (TSS 3) TSSs (**B**) of genes with high, medium, and low levels of transcription. (**C**), seven representative regions of the two-dimensional (density) signal used to characterize the interdependence on pairs of TSSs. TSSs are ordered according to their genomic position. Regions A and B correspond to transcription at the upstream TSS (0 ≤ *x* ≤ 200) when the downstream TSS is far away (−20k ≤ *y* ≤ −10k) and at an intermediate distance (−900 ≤ *y* ≤ −200), respectively. Regions Af and Bf correspond to transcription at intermediate distances from the upstream TSS (300 ≤ *x* ≤ 1*k*) when the downstream TSS is far away (−20k ≤ *y* ≤ −10k) and at an intermediate distance (−900 ≤ *y* ≤ −200), respectively. Regions C, D, and E correspond to transcription at the downstream TSS (0 ≤ *y* ≤ 200) when the upstream TSS is nearby (0 ≤ *x* ≤ 200), at an intermediate distance (300 ≤ *x* ≤ 1k), and far away (10k ≤ *x* ≤ 20k), respectively. For the quantification of proximal, intermediate, and distal effects between TSSs, we define the average transcription *T*_*W*_ in a given region *W* as 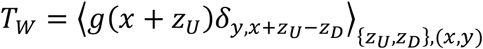 with (*x*, *y*) ∈ *W* (see Materials and Methods). Selecting *W* as one of the representative regions leads to the definitions of proximal cooperativity as *T*_*C*_/*T*_*E*_; upstream effects as *T*_*B*_/*T*_*A*_; downstream effects as *T*_*D*_/*T*_*E*_; positional dominance as *T*_*E*_/*T*_*A*_; persistence with a distal downstream TSS as *T*_*Af*_/*T*_*A*_; persistence with a non-distal downstream TSS as *T*_*Bf*_/*T*_*B*_; and signal dominance as *T*_*Bf*_/*T*_*Af*_. Data is available from the ENCODE consortium (experiment accession number ENCSR000AEL, Thomas Gingeras lab, CSHL). The accession numbers of the minus and plus strand RNA-seq signals and gene quantifications are ENCFF652ZSN, ENCFF091RAW, and ENCFF782PCD, respectively.

Many key features, however, are strongly dependent on the intragenic position of the TSSs and the transcriptional activity of the gene, which we have stratified as high, medium-high, medium, medium-low, low, and zero (Figure S2). Without this stratification, the signal would be dominated by highly transcribed genes. The most salient general feature is the absence of substantial transcription at the first annotated TSS of highly transcribed genes irrespective of its distance to the second one. Transcription at the first annotated TSS becomes more prominent only as the activity of the gene decreases. Another general salient feature is the high RNA-seq signal density just downstream of two TSSs that are close to each other.

To accurately characterize the observed dependence patterns, we consider seven regions of the two-dimensional signal density (Figure 2C). Five of the regions are located immediately downstream of one of the TSSs and are distinguished by the relative position of the other TSS, located upstream at distal and at intermediate distances (regions A and B, respectively) or downstream at proximal, at intermediate, and at distal distances (regions C, D, and E, respectively). The other two regions are located at intermediate distances downstream a TSS and at distal and at non-distal distances upstream of the next TSS (regions Af and Bf, respectively).

Explicitly, comparing RNA-seq densities in region B with those of region A indicates that the proximity of the 2nd TSS strongly correlates with reduced transcription at the 1st TSS. These *upstream effects* of the 2nd annotated TSS extend up to ~1kbp distances. In contrast, transcription in region D is higher than in region E, which shows that the *downstream effects* of the 1st annotated TSS statistically enhance transcription at the 2nd TSS. This effect is even more marked when comparing transcription in region C with transcription in region E, which we have termed *proximal cooperativity*, indicating that on average there is more transcription at the 2nd TSS the closer it is to the 1st TSS. To compare transcription when the two TSSs are far from each other, we consider regions A and E. For highly transcribed genes, transcription is much more prominent at the 2nd than at the 1st TSS. This *distal positional dominance* of the downstream TSS shifts to the upstream TSS as the transcriptional activity of the gene decreases.

The statistical interdependence of the RNA-seq signal at the first pair of annotated TSSs is also present to a large extent at the second and third TSSs (Figure 2B). Proximal, intermediate, and distal effects, except for the intermediate upstream effects for high transcription, closely parallel those of the first pair of annotated TSSs.

After transcription initiation, the average RNA-seq signal is expected to be lost progressively due to multiple processes, including transcription abortion, transcription termination, and RNA processing^23,24^. The persistence of the RNA-seq signal is strongly influenced by the position of the downstream TSS (Figures 2 and S1). Explicitly, *persistence with a non-distal downstream TSS* (average signal in region Bf compared to that of region B) is substantially higher than *persistence with a distal downstream TSS* (average signal in region Af compared to that region A), especially for low and medium values of the transcriptional activity. Therefore, the presence of a nearby downstream TSS correlates with lower transcription initiation but, at the same time, with more persistent RNA-seq signals. Regarding the absolute value of the average RNA-seq signal between TSSs, it tends to be higher as the downstream TSS gets closer to the upstream TSS (average signal in region Bf compared to that of region Af), which we refer to as *signal dominance* (Figures 2 and S1).

### Transcription initiation is statistically dependent on neighboring TSSs

To investigate the statistical interdependence of transcription initiation at neighboring annotated TSSs, we computed the two-dimensional signal densities for RNA Annotation and Mapping of Promoters for the Analysis of Gene Expression (RAMPAGE) data^25^ in the same way as for RNA-seq data (Figure 3). This technique provides specific sequencing of 5′-complete complementary DNAs and avoids counting transcripts that initiate at other TSSs. The results show that the interdependence of RAMPAGE densities at the TSSs mimics to a large extent the phenomenology observed for RNA-seq densities immediately downstream of the TSSs (Figure 2), including proximal, intermediate, and distal effects. Outside the TSS region, RAMPAGE densities are zero. The qualitative similarities between transcription initiation and transcription immediately downstream of the TSSs are consistent with a hierarchy of alternative TSS usage in delineating the overall RNA-seq signal but there are general trends in the two-dimensional RNA-seq signal density space, such as differential persistence depending on the position of the closest downstream TSS, that extend beyond transcription initiation.

**Figure 3:**
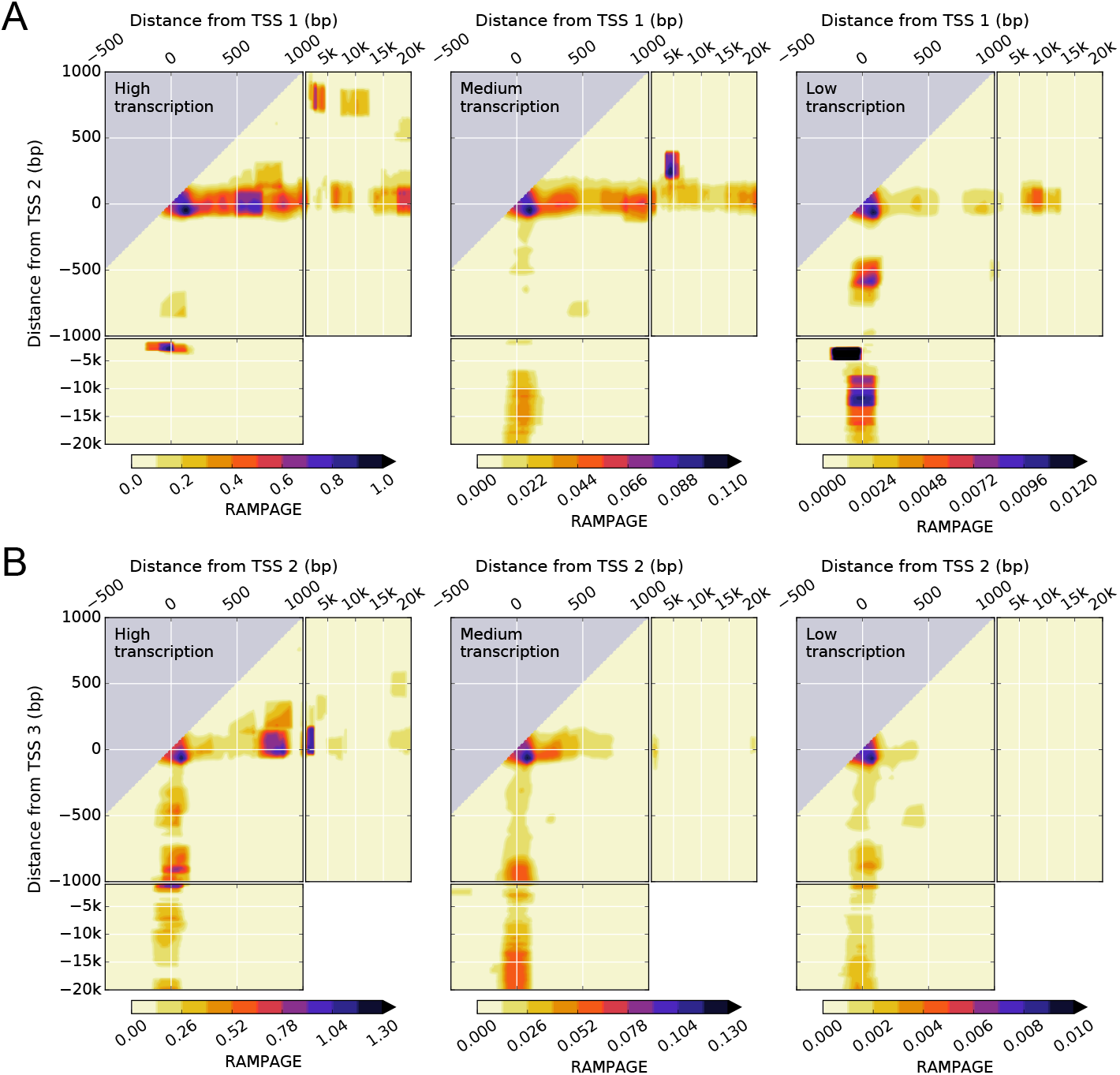
Transcription initiation in K562 leukemia cell line shows a complex dependence on the distance from pairs of TSSs, their intragenic position, and the transcriptional activity of the gene. (**A**, **B**), two-dimensional density of RAMPAGE signal for pairs of the first (TSS 1) and second (TSS 2) TSSs (**A**) and the second (TSS 2) and third (TSS 3) TSSs (**B**) of genes with high, medium, and low levels of transcription. Data is available from the ENCODE consortium (experiment accession number ENCSR000AER, Thomas Gingeras lab, CSHL). The accession numbers of the minus and plus strand RAMPAGE signals and gene quantifications are ENCFF198YEH, ENCFF707TAV, and ENCFF782PCD, respectively.

### Interdependence of transcription on neighboring TSSs is regulated

We investigated how the statistical interdependence of the RNA-seq signal on consecutive pairs of TSSs is associated with known transcriptional regulation features. Explicitly, we considered chromatin immunoprecipitation followed by sequencing (ChIP-seq) data for POLR2A as a reporter of RNA polymerase II (Pol II) occupancy (Figures 4A and S3A); DNase I hypersensitivity analysis followed by sequencing (DNase-seq) data as a reporter of DNA accessibility (Figures 4B and S3B), which is required for transcription factors and other regulatory proteins to bind DNA; and ChIP-seq data for the active chromatin marker H3K4me3 (Figures 4C and Figure S3C). The results show that changes in the transcriptional activity downstream of a TSS are correlated with changes in the transcriptional regulation features, indicating that the interdependence of transcription initiation and transcription at neighboring TSSs originates at the regulatory level.

**Figure 4:**
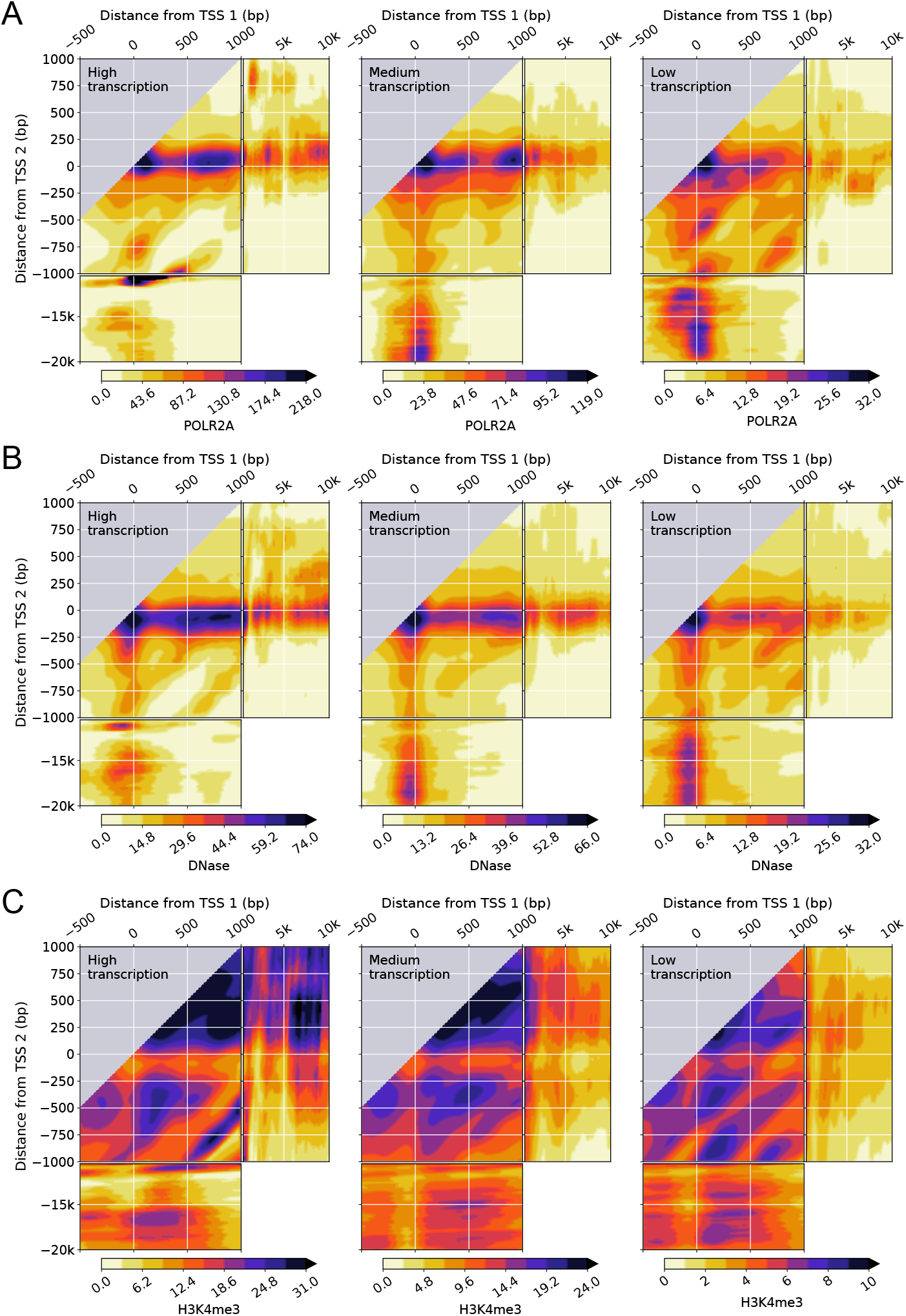
RNA polymerase II occupancy, DNA accessibility, and H3K4me3 epigenetic chemical modification of the histone H3 protein in K562 leukemia cell lines shows a complex dependence on the distance from pairs of TSSs and the transcriptional activity of the gene. (**A**, **B**, **C**), two-dimensional density of POLR2A ChIP-seq signal (**A**), DNase-seq signal (**B**), H3K4me3 ChIP-seq signal (**C**) for pairs of the first (TSS 1) and second (TSS 2) TSSs of genes with high, medium, and low levels of transcription. Data is available from the ENCODE consortium (experiment accession numbers ENCSR000FAJ, Sherman Weissman lab, Yale; ENCSR000EKS, Gregory Crawford lab, Duke; ENCSR000AKU and Bradley Bernstein, Broad). The accession numbers of the POLR2A ChIP-seq signal, DNase-seq signal, H3K4me3 ChIP-seq signal, and gene quantifications are ENCFF000YWY, ENCFF000SVL, ENCFF000BYB, and ENCFF782PCD, respectively.

Explicitly, the presence of a downstream TSS negatively impacts both Pol II occupancy and DNA accessibility around TSSs positioned upstream at intermediate distances. As in the case of transcription, these effects are much more marked for the first pair (Figures 4A and 4B) than for the second pair of TSSs (Figures S3A and S3B). Pol II occupancy and DNA accessibility are systematically enhanced as well by the presence of an upstream TSS at intermediate distances and by the cooperative actions of two proximal TSSs. For pairs of TSSs that are far from each other, the relative contributions of Pol II occupancy and DNA accessibility decrease at the downstream TSS and increase at the upstream TSS as the transcriptional activity of the gene decreases, paralleling the shift in positional dominance observed for transcription. Outside the transcription initiation regions, the RNA-seq signal closely follows the main trends of Pol II occupancy, which overlap to a large extent with DNA accessibility. Therefore, a downstream TSS affects not only transcription initiation but also transcription progression.

Pol II occupancy and the presence of DNase I hypersensitivity sites are two general indicators of transcription and of transcription initiation and regulation, respectively^26^. Similarly, the active chromatin marker H3K4me3 (Figures 4C and Figure S3C) shows differentiated patterns on the two-dimensional RNA-seq signal densities that are consistent with active transcription initiation. Namely, H3K4me3 is high downstream of transcription initiation and significantly lower just upstream. This pattern is clearly observed, for instance, for the first pair of distal TSSs around the 2nd TSS for high transcription and how it switches to the 1st TSS as transcription decreases. In general, we observe that, for high transcriptional activity, H3K4me3 is high downstream of two TSSs, low immediately upstream of both TSSs, and changing from low to high between the two TSSs depending on their relative positions.

### The interdependence of transcription on neighboring TSSs is conserved across human cell types

To study to what extent there are general trends present in other cell types, we obtained the two-dimensional RNA-seq signal densities for the GM12878 human lymphoblastoid cell line (Figure S4) and for H1-hESC human embryonic stem cells (Figure S5), which together with K562 constitute the three Tier 1 cell types of the encyclopedia of DNA elements (ENCODE) project^27-29^. The main features, involving proximal cooperativity, upstream effects, downstream effects, and positional dominance, are very similar for all three cell types.

We quantified the presence of these general trends across all the spectrum of different cell types for each of the pairs of consecutive TSSs up to the 11th TSSs in all human experiments in the ENCODE project with high replicate concordance (Table S1). These included 191 experiments with 122 different cell types (biosamples), covering all different biosample types. The results show that the complex interdependence of the transcriptional signal at multiple TSSs observed in K562, GM12878, and H1-hESC cells is conserved across all variety of human cell types (Figure 5).

**Figure 5:**
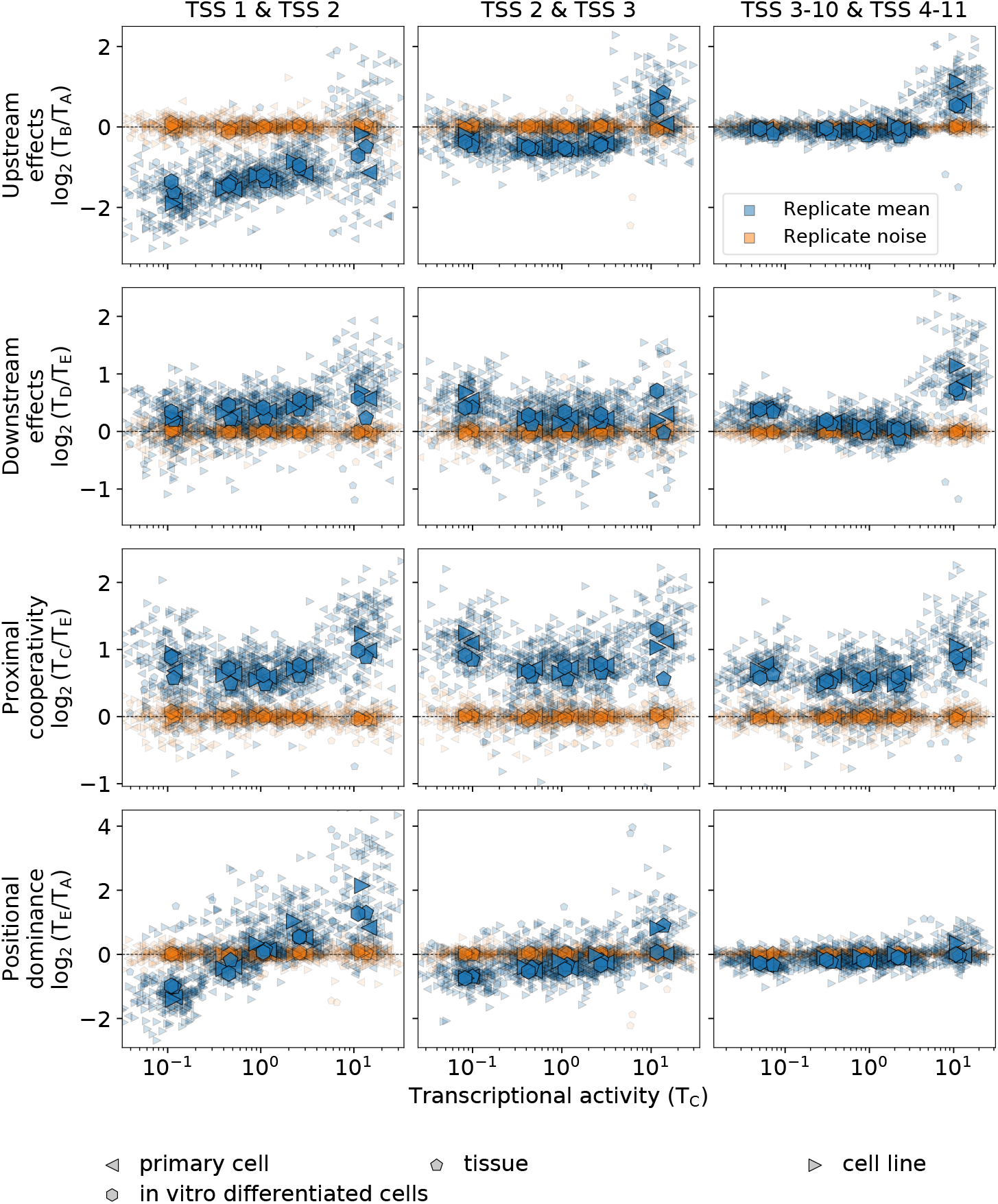
The complex interdependence of transcription at multiple TSSs is conserved across human cell types. The replicate mean and noise of the log_2_ values of *upstream effects*, *downstream effects*, *proximal cooperativity*, and *positional dominance* are shown in terms of the transcriptional activity in region C stratified in five groups for the first and second TSSs, for the second and third TSSs, and for the average of all subsequent pairs of consecutive TSS up to the 10th and 11th TSSs for all experiments in ENCODE with Spearman correlation > 0.8 among replicates. In total, there are 191 experiments (indicated by small symbols) comprising 122 different cell types. Different symbols indicate different biosample types, which include primary cell (62 experiments), cell line (93 experiments), tissue (27 experiments), and in vitro differentiated cells (9 experiments). Large symbols indicate the average of experiments within a biosample type. The replicate mean, represented in blue color, corresponds to the average of the log_2_ values of two replicates [i.e., 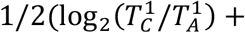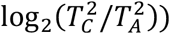, where the superscript indicates the replicate number]. The replicate noise, represented in orange color, corresponds the difference of the log_2_ value of replicate 1 from the replicate mean [i.e., 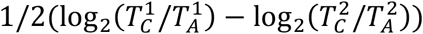]. Data is available from the ENCODE consortium (Brenton Graveley lab, UConn; Eric Lécuyer lab, IRCM; Michael Snyder lab, Stanford; and Thomas Gingeras lab, CSHL). For ENCODE accession numbers, see Table S1.

Explicitly, upstream and downstream effects are extremely marked for the first pair of TSSs, substantially decrease for the second pair, and are highly suppressed for the other pairs further downstream in the gene, except for highly transcribed genes. In this latter case, the presence of an additional TSS nearby, either upstream or downstream, is always associated with enhanced transcription. Similarly, positional dominance also ranges from very marked for the first pair of TSSs to highly suppressed for the other pairs further downstream in the gene. In contrast, proximal cooperativity is always maintained at a high level irrespective of the transcriptional activity of the gene and the relative position of the TSS pair within the gene.

We also quantified the presence of general trends in transcription initiation using RAMPAGE data for each of the pairs of consecutive TSSs up to the 11th TSSs in all human experiments in the ENCODE project with high replicate concordance (Table S2). These included 65 experiments with 56 different cell types, covering all biosample types. The results show that the main trends observed for the transcription initiation in K562 are conserved across human cell types (Figure 6). There are broad similarities with RNA-seq data but also notable differences.

**Figure 6:**
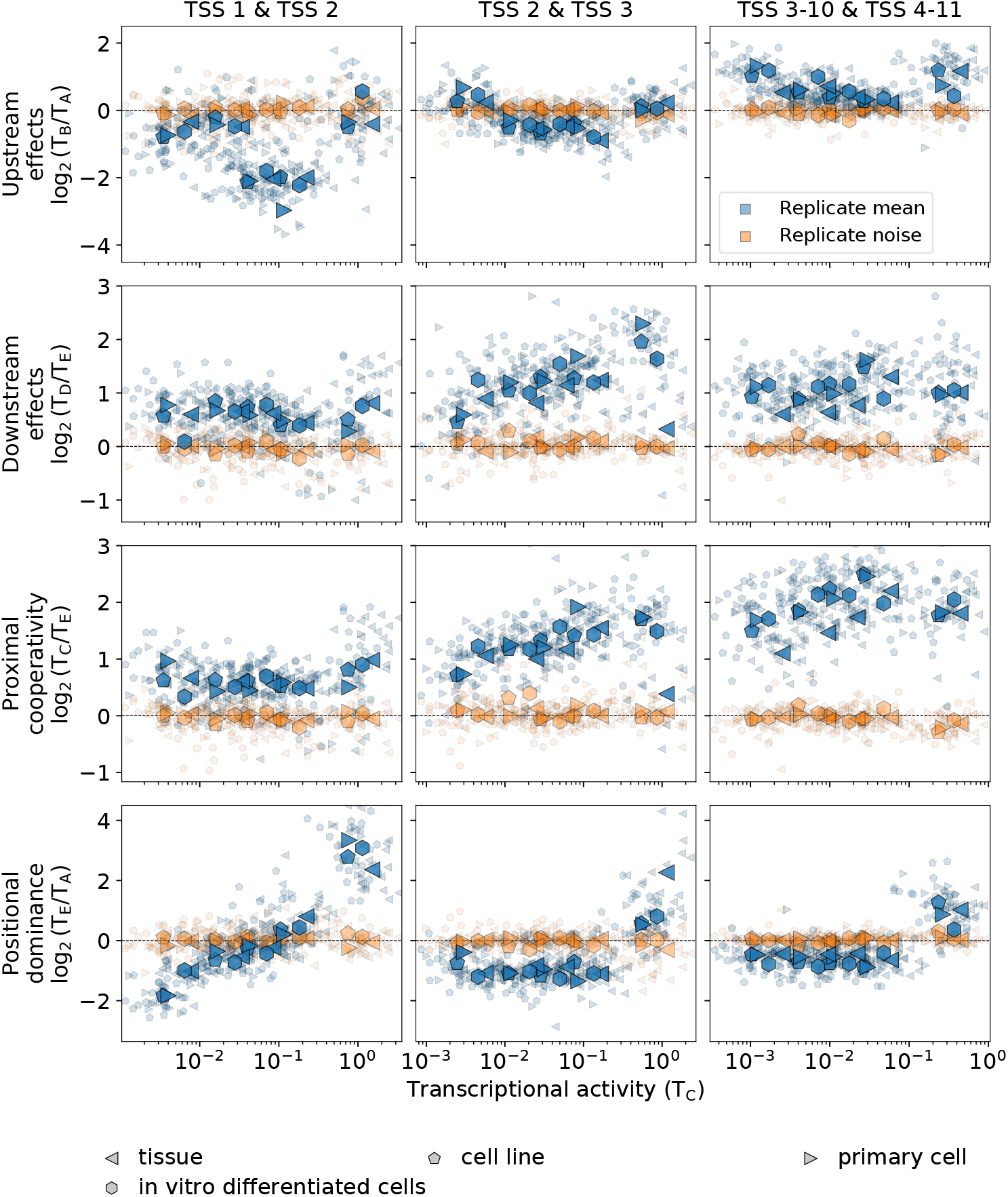
Transcription initiation parallels the conserved interdependence patterns of transcription at multiple TSSs. The same quantities as in Fig. 5 are shown computed with RAMPAGE data instead of with RNA-seq data. In total, there are 65 experiments comprising 56 different cell types, which include, as biosample types, primary cell (11 experiments), cell line (25 experiments), tissue (24 experiments), and in vitro differentiated cells (5 experiments). Data is available from the ENCODE consortium (Thomas Gingeras lab, CSHL). For ENCODE accession numbers, see Table S2.

Positional dominance for RAMPAGE data across multiple cell types closely mimics the results for RNA-seq data, indicating that transcription initiation as well as transcription generally shift from the upstream to the downstream TSS of the distal TSS pair as the transcriptional activity of the gene increases. Upstream effects are also remarkably similar for both processes, except for the first pair of TSSs with low transcriptional activity of the gene. Downstream effects and proximal cooperativity are positive in both RNA-seq and RAMPAGE data but are much more marked in the latter. In general, it is observed that these effects become more pronounced in RAMPAGE data as the positional order of the TSS pair in the gene increases. The fact that these marked transcription initiation effects are reduced to a large extent in transcription as the positional order of the TSS pair increases is consistent with transcription at a given position accounting for the cumulative effects of transcription initiated at the upstream TSSs.

The quantification of the average RNA-seq signal between consecutive TSSs indicates that the general trends observed qualitatively for Tier 1 cell types are indeed conserved across all variety of human cell types (Figure 7). Explicitly, the expected reduced average signal after transcription initiation is observed for any location of the downstream TSSs (Figure 7), except for the first TSS pair with a non-distal downstream TSS. In the case of K562 leukemia cell line, which we analyzed explicitly at the level of the regulatory features, this behavior is also present at the level of the Pol II occupancy (Figure 4), thus indicating that it is a general feature of transcription itself. Comparing the effects of the downstream TSS location, the persistence of the average RNA-seq signal is systematically higher for a non-distal than for a distal downstream TSS for all TSS pairs (Figure 7), which we refer to as *persistence dominance*. In absolute terms, the average RNA-seq signal between TSSs does not depend on the downstream TSS distance for the first pair of TSSs of the gene, but tends to be higher in the presence of a non-distal downstream TSS for subsequent TSS pairs in the gene (Figure 7).

**Figure 7:**
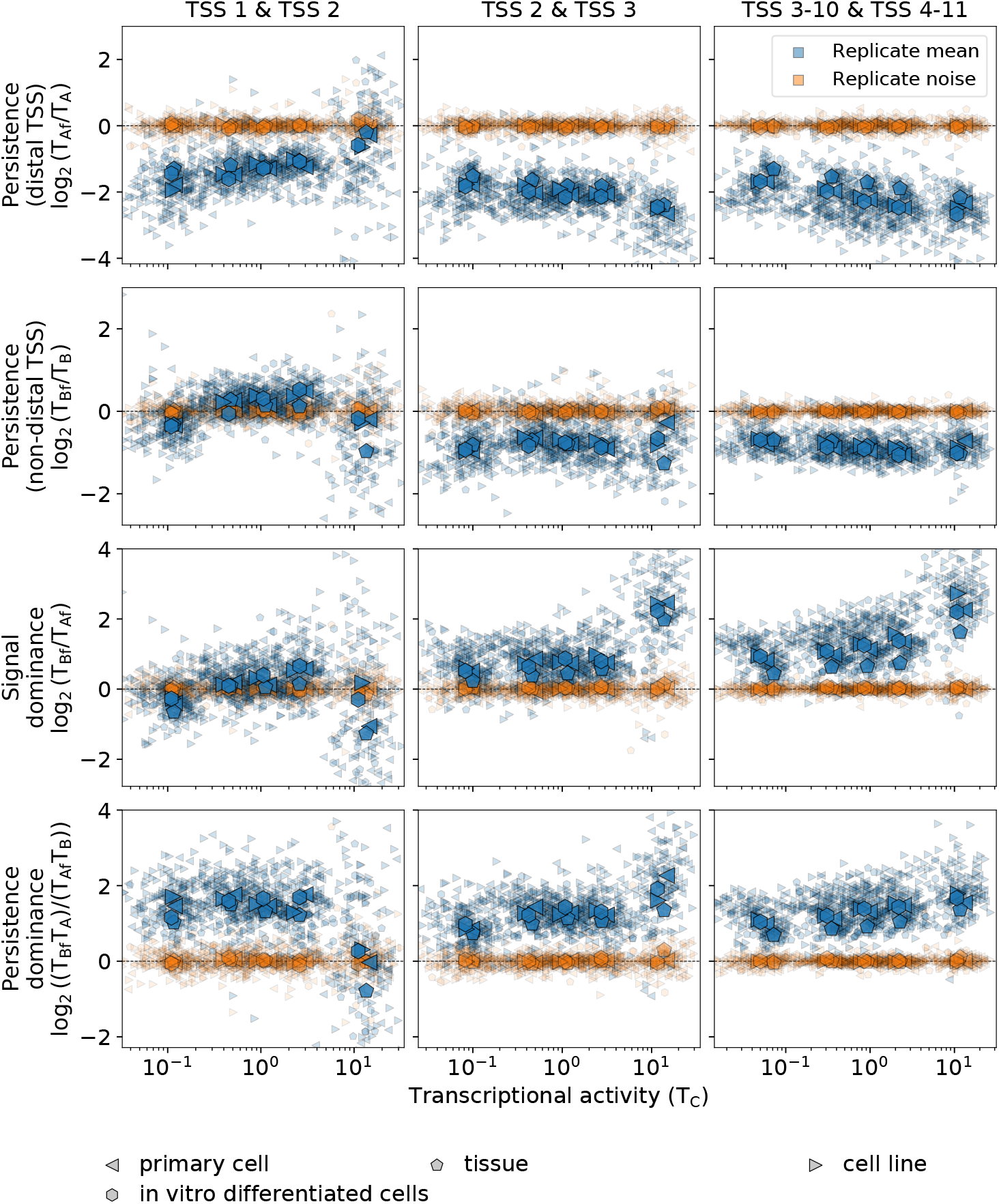
The complex interdependence of transcription between multiple TSSs is conserved across human cell types. The replicate mean and noise of the log_2_ values of transcription *persistence with a distal downstream TSS*, *persistence with a non-distal downstream TSS, signal dominance*, and *persistence dominance* are shown in terms of the transcriptional activity in region C for the same cases and conditions as in Fig. 5.

### Multiple levels of variability

Alternative transcription has important implications for gene expression as it determines the variability of the repertoire of isoform proteins. We observe that, along the main trends, there is high variability in the replicate means that is substantially higher than the variability between replicates (Figures S6, S7, and S8). The variability has both a random-like component and a bias determined by the genomic context. The bias, resulting from the general interdependence patterns across TSSs we have uncovered, is strongly emphasized in the averages of all experiments within each biosample type, which exhibit little variation across different biosample types (Figures 5, 6, and 7). Therefore, the main trends in the interdependence of transcriptional processes on the TSSs arrangements are present in the same form for all cell types, irrespective of their mutational background, specific origin, and function within the organism. Compounded with these general trends, there are multiple levels of variability, such as replicate noise, cell-type-specific TSS usage, and specific responses to different conditions.

## Discussion

Directed analyses on specific systems have shown that many fundamental mechanisms involved in transcription regulation strongly depend on the precise distances among the locations of multiple DNA elements^15,30^ but it has been unclear to what extent this dependence could be present along the genome after the confluence of many of these, potentially opposing mechanisms^31^. Especially relevant is the case of alternative transcription^32^. There is ample evidence that multiple TSSs in most genes have independent cell-type-specific expression profiles^21^. These profiles have been found to be connected to disease states, including alternative transcription initiation at multiple TSSs that is deregulated across cancer types and patients^33^ and that exhibits well-defined, specific signatures in type 2 diabetes^34^. The types of regulation comprise a wide range of modalities, including the TSSs of a gene being coregulated, namely increasing or decreasing their expression proportionally^35^, and, on the opposite side, switching expression from one TSS to another^36^.

The multiple-landmark-alignment methodology we have developed provides an avenue to elucidate how the precise positioning of multiple landmarks reflects in DNA-dependent processes on a genome-wide scale. The simultaneous consideration of multiple distances (stratified as proximal, intermediate, and distal) has been a fundamental element of our approach to uncover the existence of regulated interdependence patterns of gene expression at alternative TSSs and between TSSs across human cell lines, primary cells, in vitro differentiated cells, and tissues. This interdependence comprises proximal cooperativity, upstream and downstream interactions, positional dominance, enhancement of transcription persistence, and attenuation of the transcriptional signal. In general, these effects are highly dependent on the intragenic position of the TSSs, the transcriptional activity of the gene, and the precise distances between TSSs, but at the same time, they are consistently conserved across human cell types, irrespective of their specific origin or function within the organism.

Among the most salient phenomena, there are proximal cooperativity and downstream effects, which encompass higher transcription downstream a TSS the closer it is to an upstream TSSs within the gene. This type of enhancement observed in transcription is also present, even more prominently, in transcription initiation. On the opposite side, our results show the presence of marked upstream effects, namely, the attenuation of the transcriptional signal and transcription initiation at an upstream TSS by the presence of a nearby downstream TSS. Simultaneously with the negative effects on the absolute levels of transcription, a downstream TSS positively enhances the persistence of transcription after its initiation. Concomitantly, DNA accessibility and Pol II densities show lower but more sustained profiles for a non-distal than for a distal downstream TSS. These results can be understood mechanistically considering that the assembly of the transcription initiation complex upstream a TSS interferes with transcription initiation and that the overall transcription process promotes upstream and downstream DNA accessibility.

The high variability present in all these effects at the specific experiment (specimen and condition) level agrees with multiple alternative transcription initiation being largely nonadaptive and resulting predominantly from imprecise events, as concluded by recent genome-wide analyses^37^. In more detail, fundamental biochemical principles dictate that non-specific effects, as quantified and validated in simpler gene expression prokaryotic systems, cannot generally be suppressed completely and that they are affected by regulatory processes in the usual way^38^. In this context, the existence of general patterns within multiple levels of variability we have identified is consistent with substantial non-specific transcription initiation, including both a random-like component and a bias determined by the genomic context.

Our analysis has also shown that there are clear positional dominance effects when the two TSSs are far from each other, resulting in most of transcription and transcription initiation shifting from the upstream to the downstream TSS of the distal TSS pair as the transcriptional activity of the gene increases. This effect is extremely marked for the 1st and 2nd annotated TSS of a gene and it is generally more pronounced for transcription initiation than for transcription. These types of results have also a practical side as they can be used to refine and complement TSS annotations. Explicitly, the case of positional dominance implies that the 1st annotated TSS of a gene is essentially not active, or it is not an actual TSS, if the transcriptional activity of the gene is high.

The most remarkable finding of our work is therefore the discovery of the existence of general regulated interdependence patterns of gene expression at and between alternative TSSs of protein-coding genes. We showed that these effects are conserved across cell types through a comprehensive analysis of the hundreds of human transcription and transcription initiation experiments of the ENCODE project. Compounded with these general patterns, there are multiple levels of variability, such as replicate noise, cell-type-specific TSS usage, and adaptation to different conditions. The identification of these general patterns in the alternating structure of transcription has important implications for gene expression as they determine the variability of the repertoire of isoform proteins.

On the methodological side, our approach can generally be applied to virtually any combination of landmarks and any genomic signal in the same way as we have applied them to TSSs and RNA-seq, RAMPAGE, DNase-seq, and ChIP-seq signals. Therefore, our results open an avenue to find novel distance-dependent functional relationships among multiple DNA elements in a wide variety of systems.

## Materials and methods

### Genomic signals in two dimensions

The average of the signal 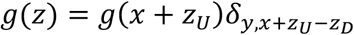 over a rectangular region from *x*_0_ to *x*_1_ along the *x* coordinate and from *y*_0_ to *y*_1_ along the *y* coordinate for all TSS pairs in the set *V* is expressed as

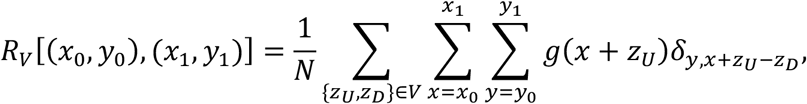

where *N* is the normalization factor, which is given by

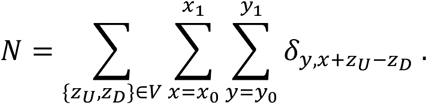

In our analysis, we use multiple sets *V* corresponding to a specific contiguous pair of TSSs of genes with transcriptional activities within a range of values (e.g., the set of 1st and 2nd TSSs of all protein-coding genes with high transcription).

### Two-dimensional region averages

To compute the region average from the previous expression efficiently, we take into account that 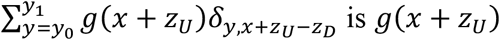 is *g*(*x* + *z*_*U*_) if *y*_0_ ≤ *x* + *z*_*U*_ − *z*_*D*_ ≤ *y*_1_ and zero otherwise. Therefore, the sum over *x* is different from zero only for *x* ≥ *y*_0_ − *z*_*U*_ + *z*_*D*_ and *x* ≤ *y*_1_ − *z*_*U*_ + *z*_*D*_, which leads to

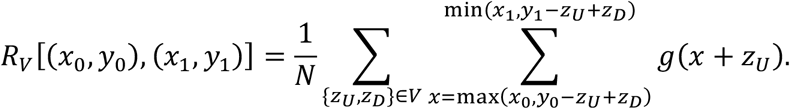

Similarly, the normalization factor is expressed as

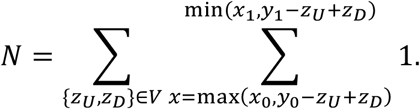

Note that the region average is defined only if there exists at least a pair {*z*_*U*_, *z*_*D*_} in *V* so that *y*_0_ − *x*_1_ ≤ *z*_*U*_ − *z*_*D*_ ≤ *y*_D_ − *x*_0_, which is equivalent to the condition max(*x*_0_, *y*_0_ − *z*_*U*_ + *z*_*D*_) ≤ min(*x*_1_, *y*_1_ − *z*_*U*_ + *z*_*D*_).

### Two-dimensional signal densities

To compute the signal densities, we use a moving window defined by a rectangular domain centered at (*x*, *y*) with dimensions 2*n*_*X*_ + 1 along the *x* coordinate and 2*n*_*Y*_ + 1 along the *y* coordinate. The signal density *G*(*x*, *y*) averaged over this domain for all TSS pairs in the set *V* is given by

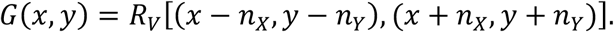

The explicit values of *n*_*X*_ and *n*_*Y*_ used in our analysis are *n*_*X*_ = 99 for −500 ≤ *x* ≤ 1*k*, *n*_*X*_ = *x*/4 for 1*k* < *x* ≤ 20*k*, *n*_*Y*_ = 99 for −1*k* ≤ *y* ≤ 1*k*, and *n*_*Y*_ = −*y*/4 for −20*k* ≤ *y* < −1*k*.

### Average transcription in a region

The average transcription in a region *W*, 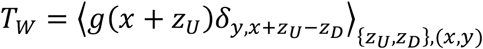 with (*x*, *y*) ∈ *W*, is computed explicitly for the representative regions as *T*_*A*_ = *R*_*V*_[(0, −20*k*), (200, −10*k*)], *T*_*Af*_ = *R*_*v*_[(300, −20*k*), (1*k*, −10*k*)], *T*_*B*_ = *R*_*v*_[(0, −900), (200, −200)], *T*_*Bf*_ = *R*_*v*_[(300, −900), (1*k*, −200)], *T*_*C*_ = *R*_*v*_[(0, 0), (200, 200)], *T*_*D*_ = *R*_*v*_[(300, 0), (1*k*, 200)], and *T*_*E*_ = *R*_*v*_[(10*k*, 0), (20*k*, 200)].

### TSSs

TSSs were obtained from the comprehensive gene annotation on the reference chromosomes of Gencode V19 (https://www.gencodegenes.org/human/release_19.html).

### TSS order

TSSs are ordered according to their genomic position, starting the enumeration from the most upstream TSS. Therefore, according to this notation, the 1st TSS does not necessarily correspond to the TSS with the highest expression.

### Genomic signals

RNA-seq, RAMPAGE, DNase-seq, and ChIP-seq genomic signals were downloaded from the Encyclopedia of DNA Elements (ENCODE) consortium repository (http://www.encodeproject.org/) as bigWig files for the hg19 mapping assembly/V19 genome annotation. Gene quantifications for the corresponding RNA-seq signals were downloaded as tsv files. Signals were normalized by their average value over the whole genome before analysis.

### RNA-seq experiment selection

RNA-seq experiments were selected in two steps. First, we considered all the experiments that matched the search criteria "hg19" for assembly and "polyA mRNA RNA-seq" or "total RNA-seq" for assay title, which produced 594 results. Subsequently, we selected RNA-seq experiments that included plus and minus strand signal of unique reads and that had high replicate concordance (Spearman correlation >0.8 between gene quantifications of the replicates), which resulted in 191 experiments.

### RAMPAGE experiment selection

RAMPAGE experiments were selected in two steps. First, we considered all the experiments that matched the search criteria "hg19" for assembly and "RAMPAGE" for assay title, which produced 155 results. Subsequently, we selected RAMPAGE experiments that included plus and minus strand signal of unique reads and that had high replicate concordance (Spearman correlation >0.8 between gene quantifications of the replicates), which resulted in 65 experiments.

## Acknowledgements

This work was supported by Ministerio de Ciencia e Innovación (MCI/AEI/FEDER, UE PGC2018-101282-B-I00 and PID2021-128850NB-I00 to J.M.G.V.) and the University of California, Davis (to L.S.).

## Author contributions

J.M.G.V and L.S conceived, designed, and performed the research.

## Supplemental information

Document S1 (Figures S1-S8, legends for tables S1 and S2); Table S1; Table S2.

## Supplemental Information

**Figure S1:**
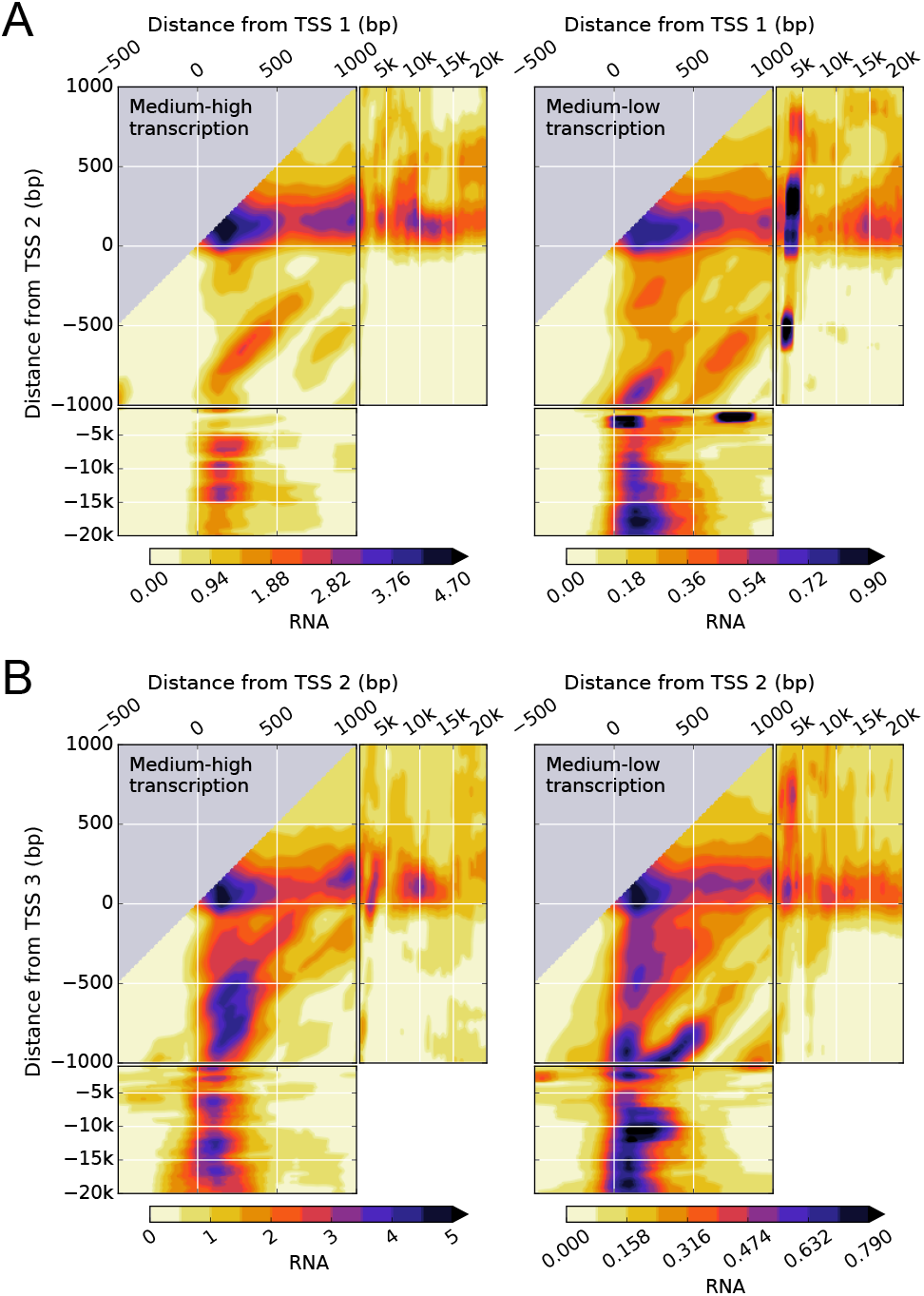
Transcription in K562 leukemia cell lines shows a complex dependence on the distance from pairs of TSSs, their intragenic position, and the transcriptional activity of the gene. (**A**, **B**), two-dimensional density of RNA-seq signal for pairs of the first (TSS 1) and second (TSS 2) TSSs (**A**) and the second (TSS 2) and third (TSS 3) TSSs (**B**) of genes with medium-high, and medium-low levels of transcription for the same conditions as in Fig. 2.

**Figure S2:**
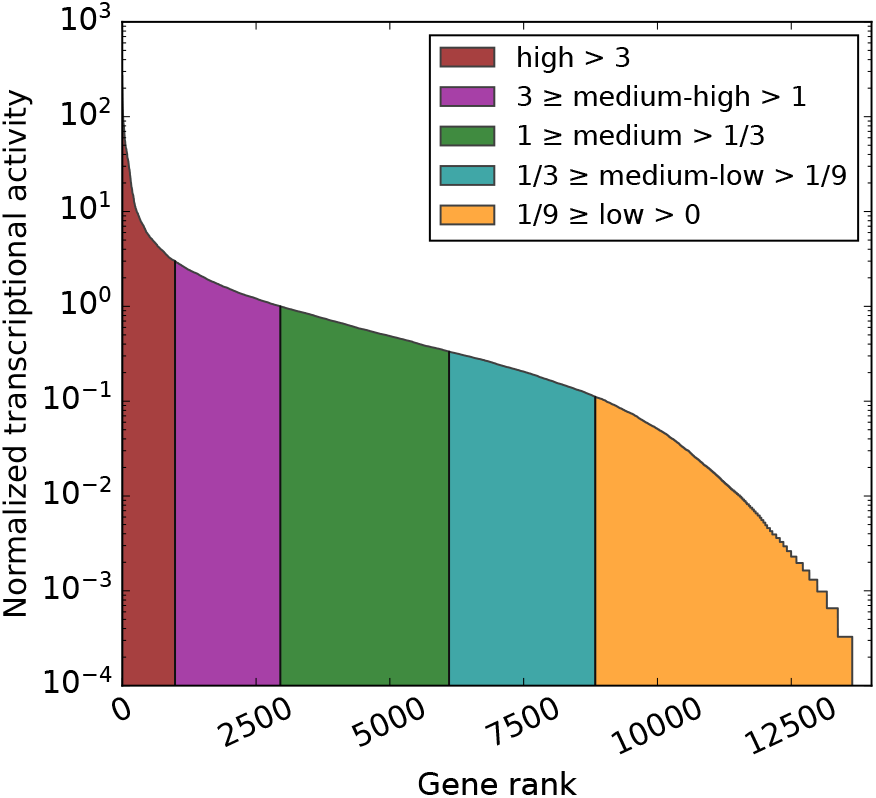
Normalized transcriptional activity (transcription / average transcription) of the protein coding genes in the K562 leukemia cell line ordered according to their transcriptional activity. The transcriptional activity of the gene was stratified as high, medium-high, medium, medium-low, and low. The number of genes in each category is 987, 1967, 3150, 2735, 4802, and 6685, respectively, of a total of 20327 protein-coding genes. Genes with zero transcriptional activity (zero RNA-seq read counts) are not considered. The ENCODE accession number for the gene quantifications is ENCFF782PCD.

**Figure S3:**
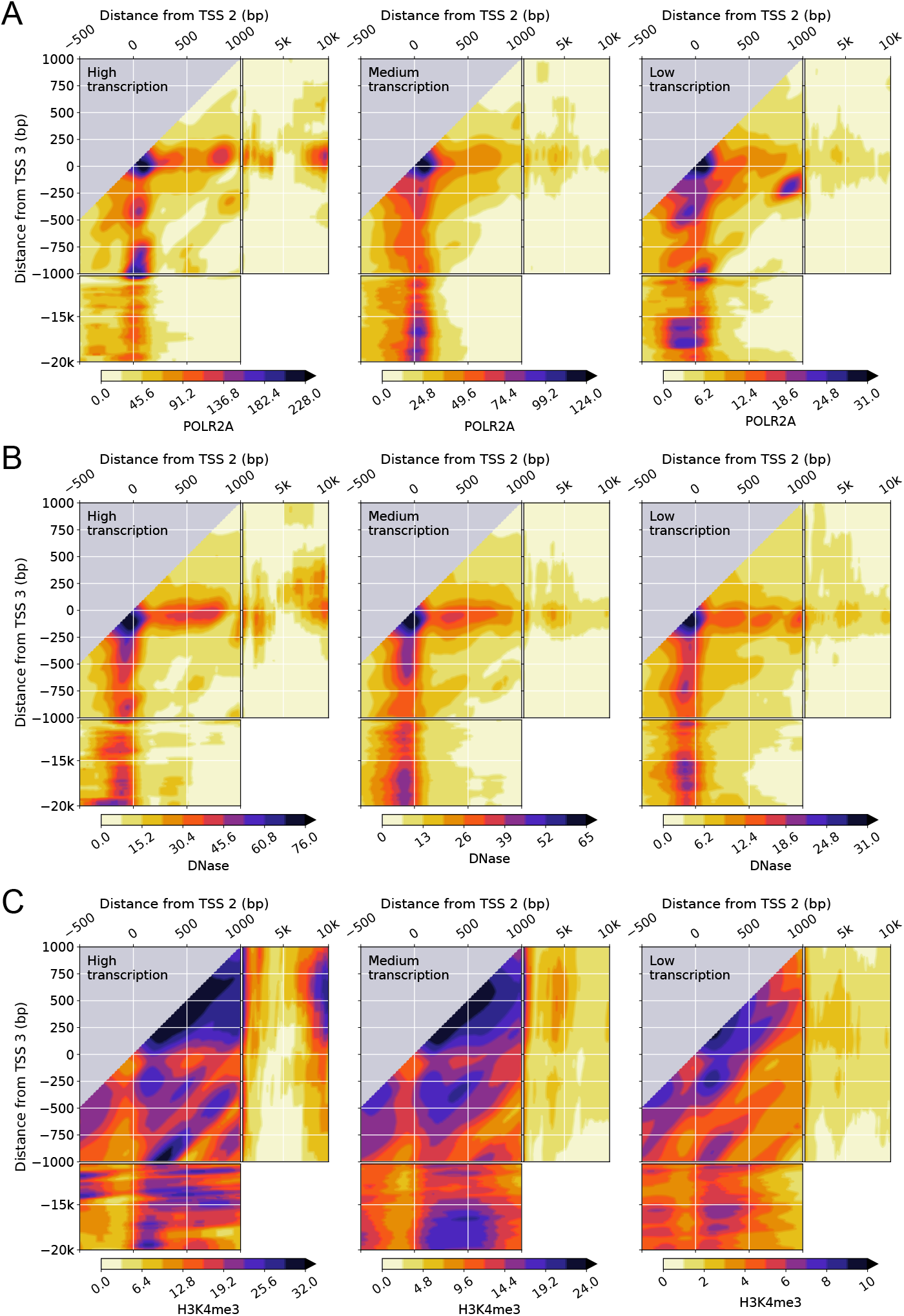
The intragenic position of the pair of TSSs affects the dependence of the RNA polymerase II occupancy, DNA accessibility, and H3K4me3 epigenetic chemical modification of the histone H3 protein on the distance from pairs of TSSs and the transcriptional activity of the gene in K562 leukemia cell lines. (**A**, **B**, **C**), two-dimensional density of POLR2A ChIP-seq signal (**A**), DNase-seq signal (**B**), H3K4me3 ChIP-seq signal (**B**) as in Fig. 4 but for pairs of the second (TSS 2) and third (TSS 3) TSSs instead of for pairs of the first (TSS 1) and second (TSS 2) TSSs.

**Figure S4:**
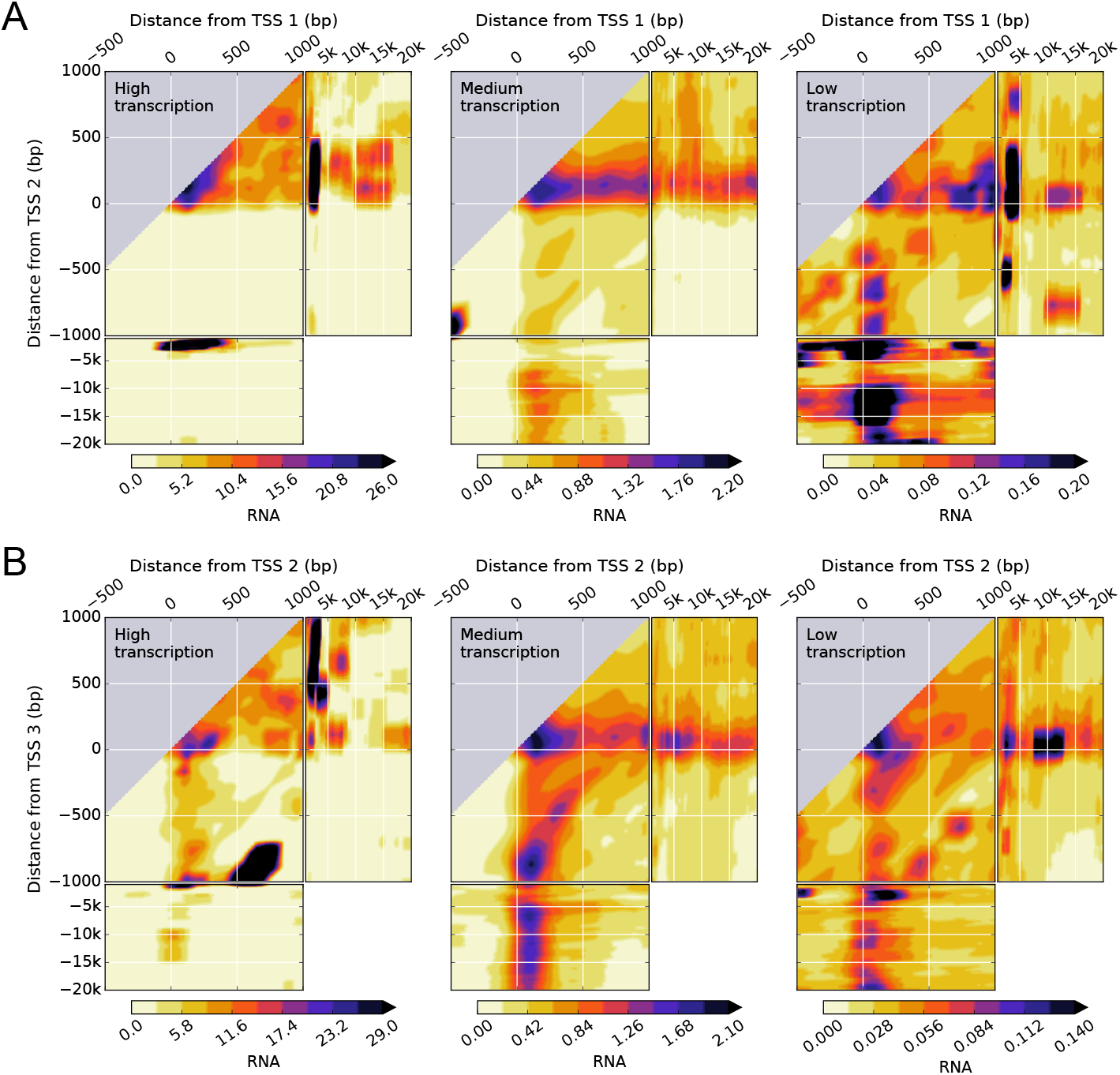
Transcription in GM12878 human lymphoblastoid cell line shows a complex dependence on the distance from pairs of TSSs, their intragenic position, and the transcriptional activity of the gene. (**A**, **B**), two-dimensional density of RNA-seq signal for pairs of the first (TSS 1) and second (TSS 2) TSSs (**A**) and the second (TSS 2) and third (TSS 3) TSSs (**B**) of genes with high, medium, and low levels of transcription. Data is available from the ENCODE consortium (experiment accession number ENCSR000AEC, Thomas Gingeras lab, CSHL). The accession numbers of the minus and plus strand RNA-seq signals and gene quantifications are ENCFF339IYH, ENCFF270HBR, and ENCFF610BLQ, respectively.

**Figure S5:**
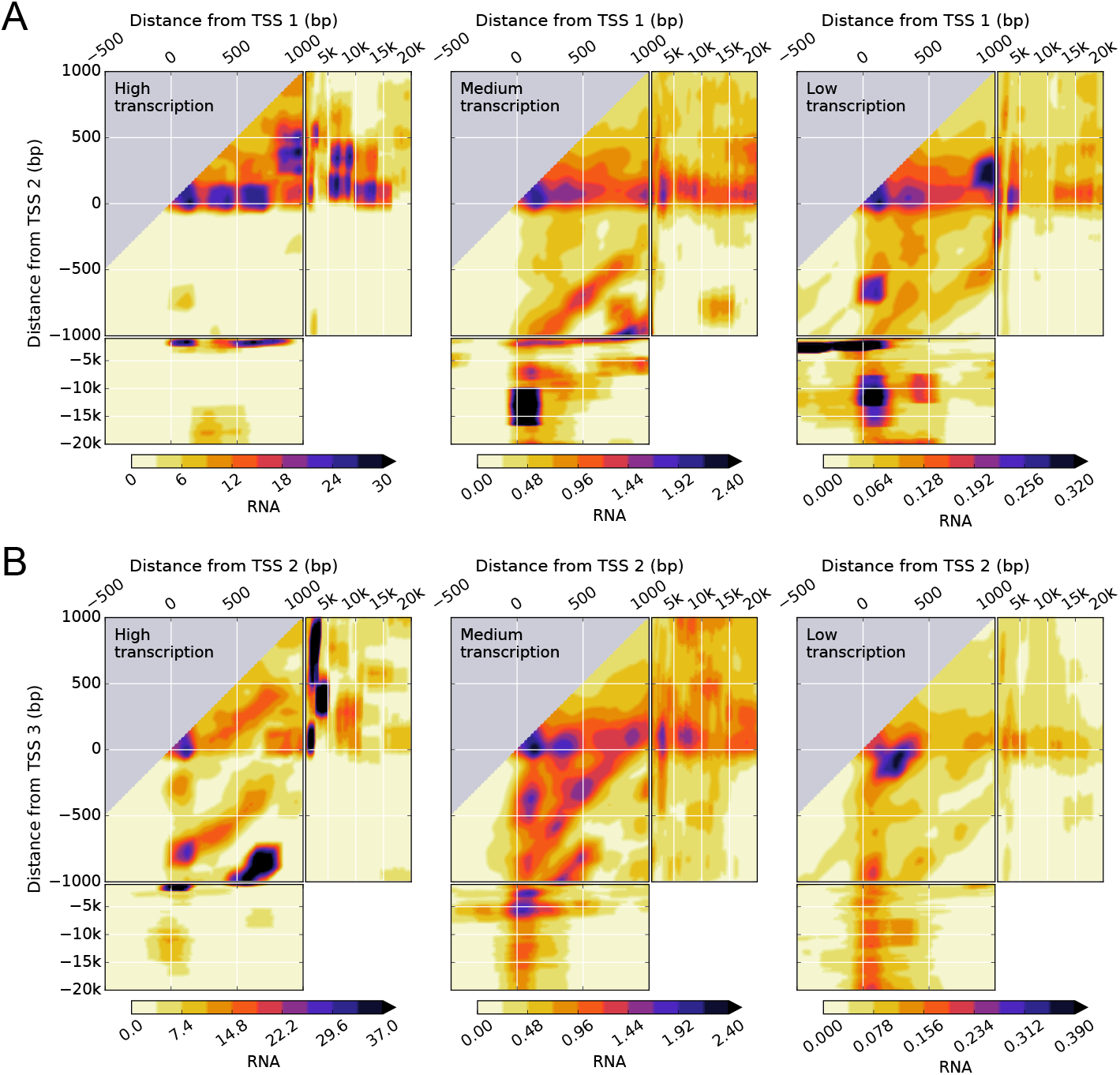
Transcription in H1-hESC human embryonic stem cells shows a complex dependence on the distance from pairs of TSSs, their intragenic position, and the transcriptional activity of the gene. (**A**, **B**), two-dimensional density of RNA-seq signal for pairs of the first (TSS 1) and second (TSS 2) TSSs (**A**) and the second (TSS 2) and third (TSS 3) TSSs (**B**) of genes with high, medium, and low levels of transcription. Data is available from the ENCODE consortium (experiment accession number ENCSR000COU, Thomas Gingeras lab, CSHL). The accession numbers of the minus and plus strand RNA-seq signals and gene quantifications are ENCFF587UUL, ENCFF587UUL, and ENCFF334LZL, respectively.

**Figure S6:**
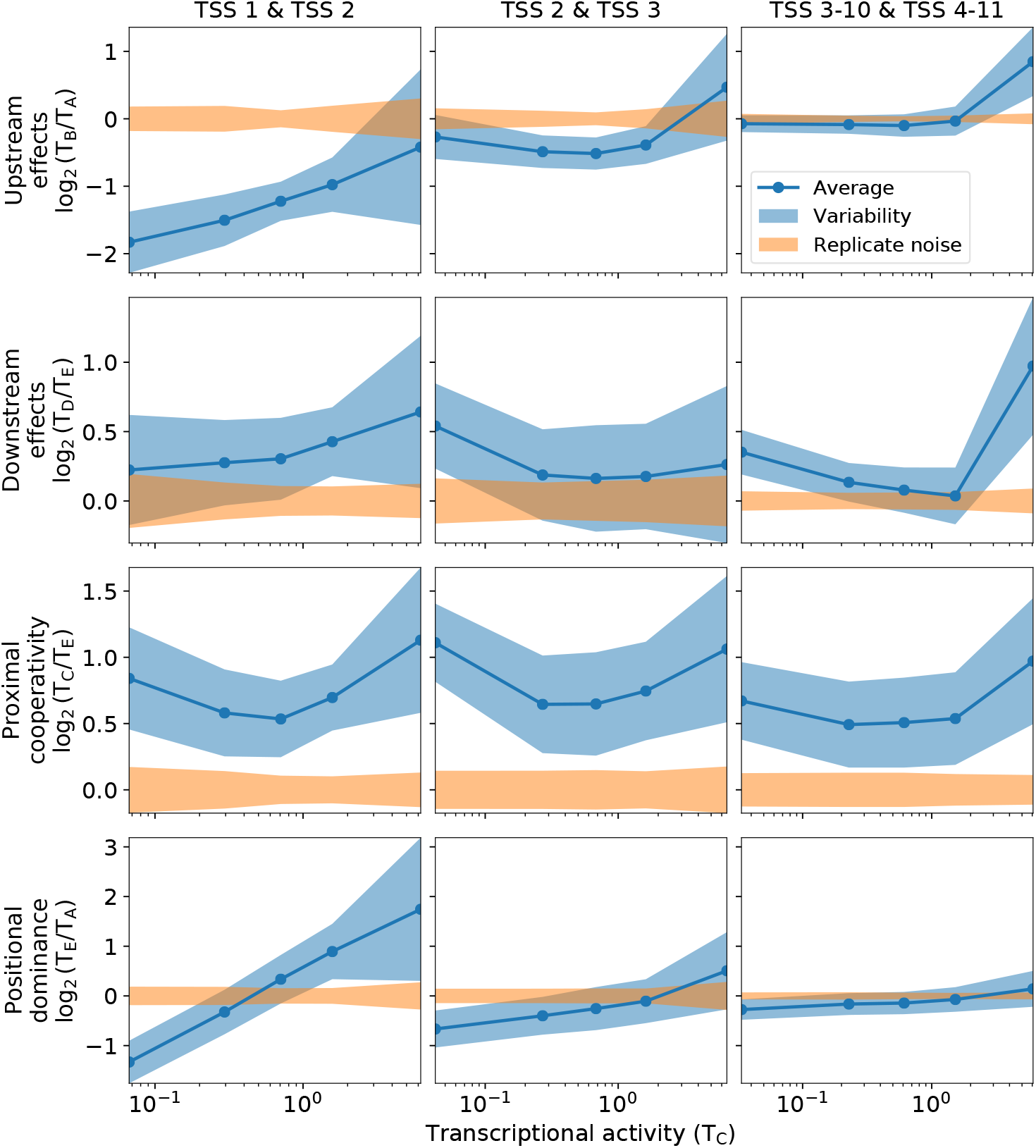
Average behavior, variability, and replicate noise of the interdependence of transcription at TSSs across human cell types. The statistical characterization of the interdependence is shown for the data of Fig. 5 for each of the five transcriptional levels. The blue lines represent the average of the log_2_ values over all the experiments; the blue-shaded region represents the variability computed as the ± standard deviation of the log_2_ replicate means of all the experiments; and the orange-shaded region represents the ± standard deviation of the replicate noise.

**Figure S7:**
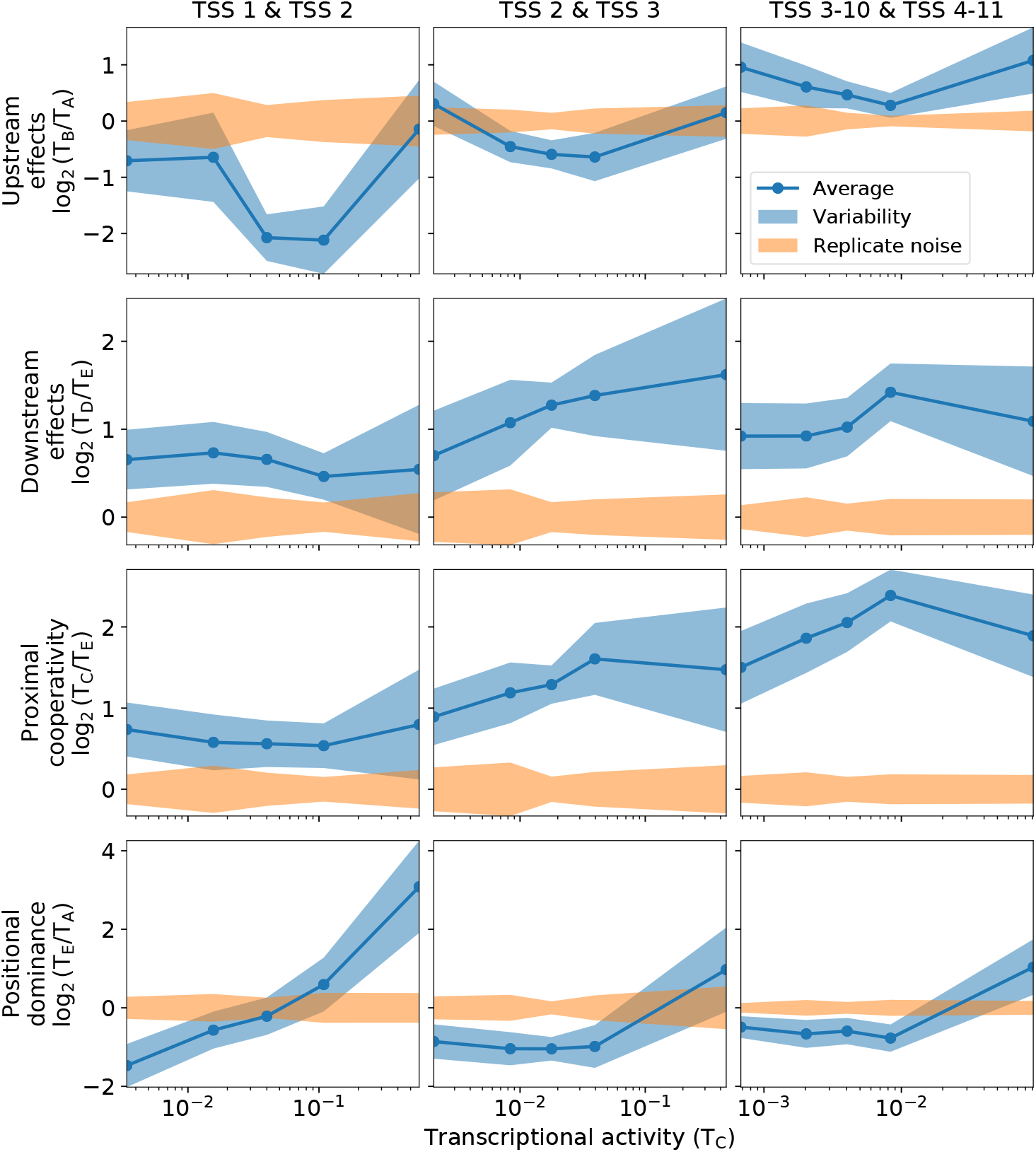
Average behavior, variability, and replicate noise of the interdependence of transcription initiation at TSSs across human cell types. The statistical characterization of the interdependence is shown for the data of Fig. 6 for each of the five transcriptional levels. The blue lines represent the average of the log_2_ values over all the experiments; the blue-shaded region represents the variability computed as the ± standard deviation of the log_2_ replicate means of all the experiments; and the orange-shaded region represents the ± standard deviation of the replicate noise.

**Figure S8:**
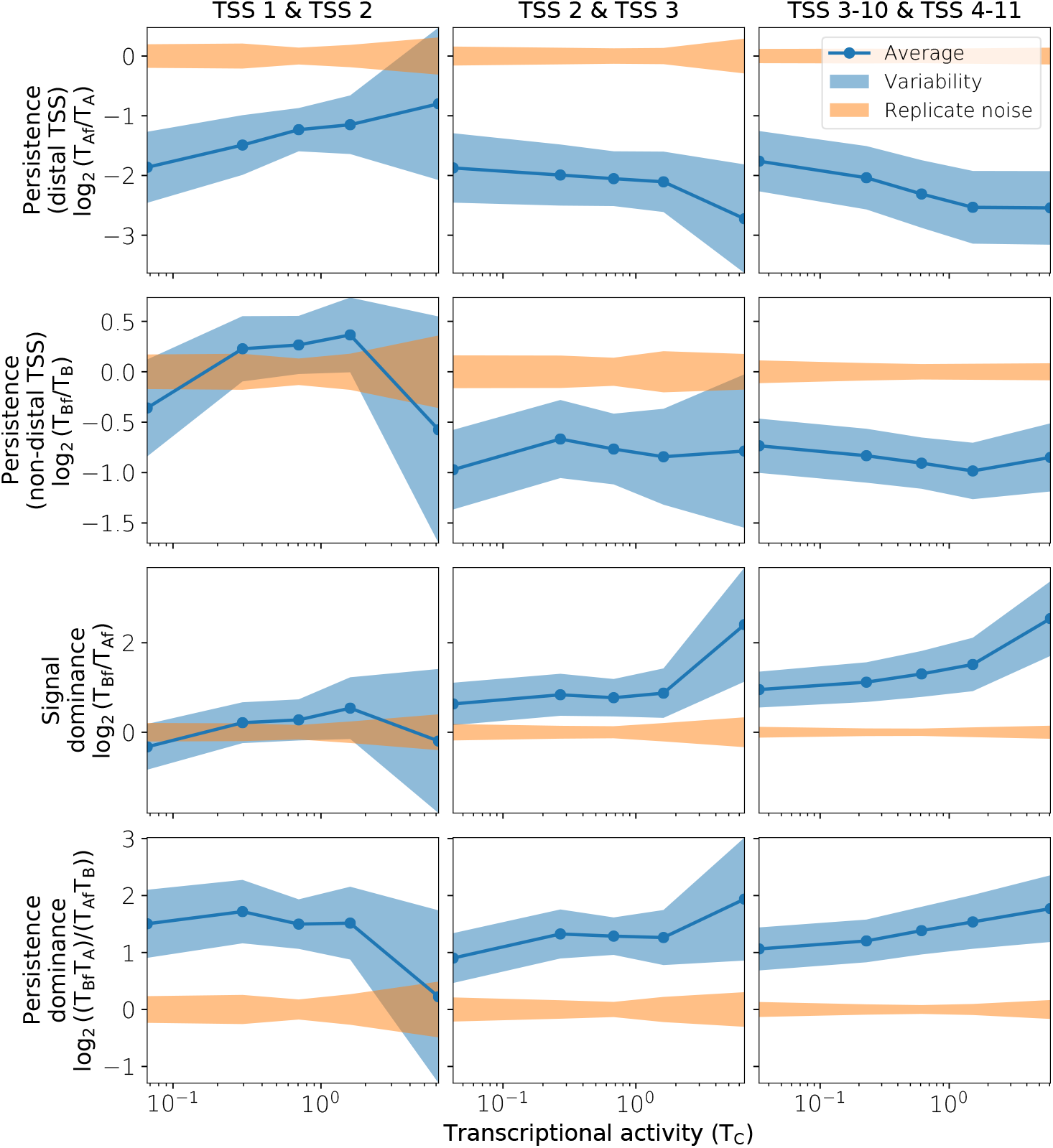
Average behavior, variability, and replicate noise of the interdependence of transcription between TSSs across human cell types. The statistical characterization of the interdependence is shown for the data of Fig. 7 for each of the five transcriptional levels. The blue lines represent the average of the log_2_ values over all the experiments; the blue-shaded region represents the variability computed as the ± standard deviation of the log_2_ replicate means of all the experiments; and the orange-shaded region represents the ± standard deviation of the replicate noise.

## Legends for supplemental tables

**Table S1.** Quantification of the average transcription in the seven representative regions of the two-dimensional signal density for different cell types for each of the pairs of consecutive TSSs up to the 11th TSSs in all human experiments in the ENCODE project with high replicate concordance. The data includes the experiment accession number, the biosample name, the biosample summary, the biosample type, the assay, the accession number for the plus strand, the accession number for the minus strand, the Spearman correlation between replicates, the replicate number, the stratified transcriptional activity level, the intragenic position of the TSS pair, and the corresponding average transcription in the regions A, Af, B, Bf, C, D, and E, labeled as T_A, T_Af, T_B, T_Bf, T_C, T_D, and T_E, respectively.

**Table S2.** Quantification of average transcription initiation performed in the same way as the quantification of average transcription in Table S1.

